# Microeconomics of metabolism: Overflow metabolism as Giffen behaviour

**DOI:** 10.1101/613166

**Authors:** Jumpei F. Yamagishi, Tetsuhiro S. Hatakeyama

## Abstract

Biological systems such as intracellular metabolism are rationally regulated to maximize the cellular growth rate through evolution (1–4), whereas microeconomics investigates the behaviour of consumers assumed to rationally maximize their utility (5, 6). Despite this analogy as optimization problems, the link between biology and economics has not been fully established (7, 8). Here, we developed an exact mapping between the regulation of metabolism and the theory of consumer choice, thereby revealing the correspondence between long-standing mysteries in both fields: overflow metabolism and Giffen behaviour. Overflow metabolism, particularly known as the Warburg effect in cancer (9, 10), is a seemingly wasteful but ubiquitous strategy where cells utilize aerobic glycolysis instead of the more energetically-efficient oxidative phosphorylation (9–16), whereas Giffen behaviour is the unexpected consumer behaviour where a good is demanded more as its price rises (17, 18). We revealed that the general conditions for these phenomena are trade-off and complementarity, i.e., impossibility of substitution for different goods. This correspondence implies that oxidative phosphorylation is counterintuitively stimulated when its efficiency is decreased by metabolic perturbations like drug administration. Therefore, Giffen behaviour bridges the Warburg effect and the reverse and inverse Warburg effect (19–22). This highlights that application of microeconomics to metabolism can offer new predictions and paradigms for both biology and economics.

---

Metabolic behaviours can be often explained as a consequence of rational regulation to optimize cellular growth rate, as successfully predicted by biological theories such as flux balance analysis (FBA) (1–4). In contrast, microeconomics describes the behaviour of individuals assumed to have perfect rationality to maximize their utility (Box 1). There is an apparent analogy between the two fields as optimization problems (7, 8). We pushed beyond this analogy to precisely map metabolism onto the theory of consumer choice in microeconomics, and examined the correspondence between overflow metabolism and Giffen behaviour.

Overflow metabolism is a seemingly wasteful behaviour, also known as the Warburg effect as a therapeutic target in cancer (21–23), where aerobic glycolysis is favoured over the more energetically-efficient oxidative phosphorylation by fast-growing mammalian cells. Those are termed fermentation and respiration in studies on microbial cells. It is ubiquitously observed even in the presence of abundant oxygen, across a variety of cells ranging from bacteria to eukaryotes such as cancer cells (9, 10), stem cells (11), immune cells (12), yeasts (13), and *E. coli* (14). In the context of the Warburg effect for cancer, many hypotheses have been proposed: e.g., respiration requires larger solvent capacity or intracellular volume because the turnover of glycolytic enzymes is much faster (15, 16); respiration retards the cellular redox state (24); glycolysis with lactate secretion is more efficient in the production of another essential metabolite, NADPH (9). Similar hypotheses have also been proposed for overflow metabolism in bacteria (16, 25–27). Nevertheless, no unified theory of this ubiquitous phenomenon has been developed. In addition, reversal of the Warburg effect, called the inverse or reverse Warburg effect, has been recently reported wherein drug administration, mitochondrial inefficiency, or intercellular interaction between cancer and stromal cells stimulates respiration (19–22). Its relationship with the Warburg effect is still unclear, and it cannot be explained by the existing hypotheses.

In microeconomics, Giffen behaviour represents the counterintuitive phenomenon where the demand for a good increases when its price rises. This is in stark contrast to the general pattern of human economic activities in which the demand for normal goods decreases when their prices rise, called Giffen’s paradox (5, 17). Even though the existence of Giffen goods was theoretically predicted more than a century ago and a few possible examples have been considered, their practical existence and mechanism remain controversial (17, 18).

Here, We apply the theory of consumer choice to understand metabolic systems (Table 1). Our general theory shows that a trade-off is essential for overflow metabolism and that all types of trade-offs in the aforementioned hypotheses can be formulated as an identical optimization problem with the same universal structure (detailed in Supplementary Note 2). In the main text, we explain only one of these for simplicity, i.e., the allocation of limited resources other than nutrients imposes a trade-off. We consider a simple metabolic system that comprises the intake flux of the carbon source as a nutrient, *J*_*C,in*_, and fluxes to metabolize the nutrient to energy molecules in the electron transport chain and TCA cycle, *J*_*C,e*_, and glycolytic pathways, *J*_*C,g*_ (Fig. 1a). *J*_*C,in*_ corresponds to the income, and *J*_*C,e*_ and *J*_*C,g*_ correspond to the demand for goods. The budget constraint line is thus given by the carbon balance as

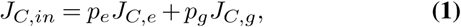

where the “prices” of the electron transport chain and aerobic glycolysis, *p*_*e*_ and *p*_*g*_, are the inverse of the efficiencies in their metabolising the nutrient to downstream metabolites. Note here that the price in economics quantifies the inefficiency of conversion from money to goods. As economic experiments control the price of goods, the price of each metabolic pathway is defined as a quantity controllable without any genetic manipulation. For example, if leakage or degradation of intermediate metabolites is increased by drugs, the price of the metabolic pathways increases.

**Table 1.** Mapping between microeconomics and metabolism

| Microeconomics | Metabolism |
| --- | --- |
| Utility | Growth rate |
| Income | Intake of nutrient |
| Goods | Metabolic pathways |
| Demand for goods | Allocation of nutrient |
| Price of goods | Inefficiency of metabolism |
| Complementarity | Stoichiometry |

**Fig. 1.**
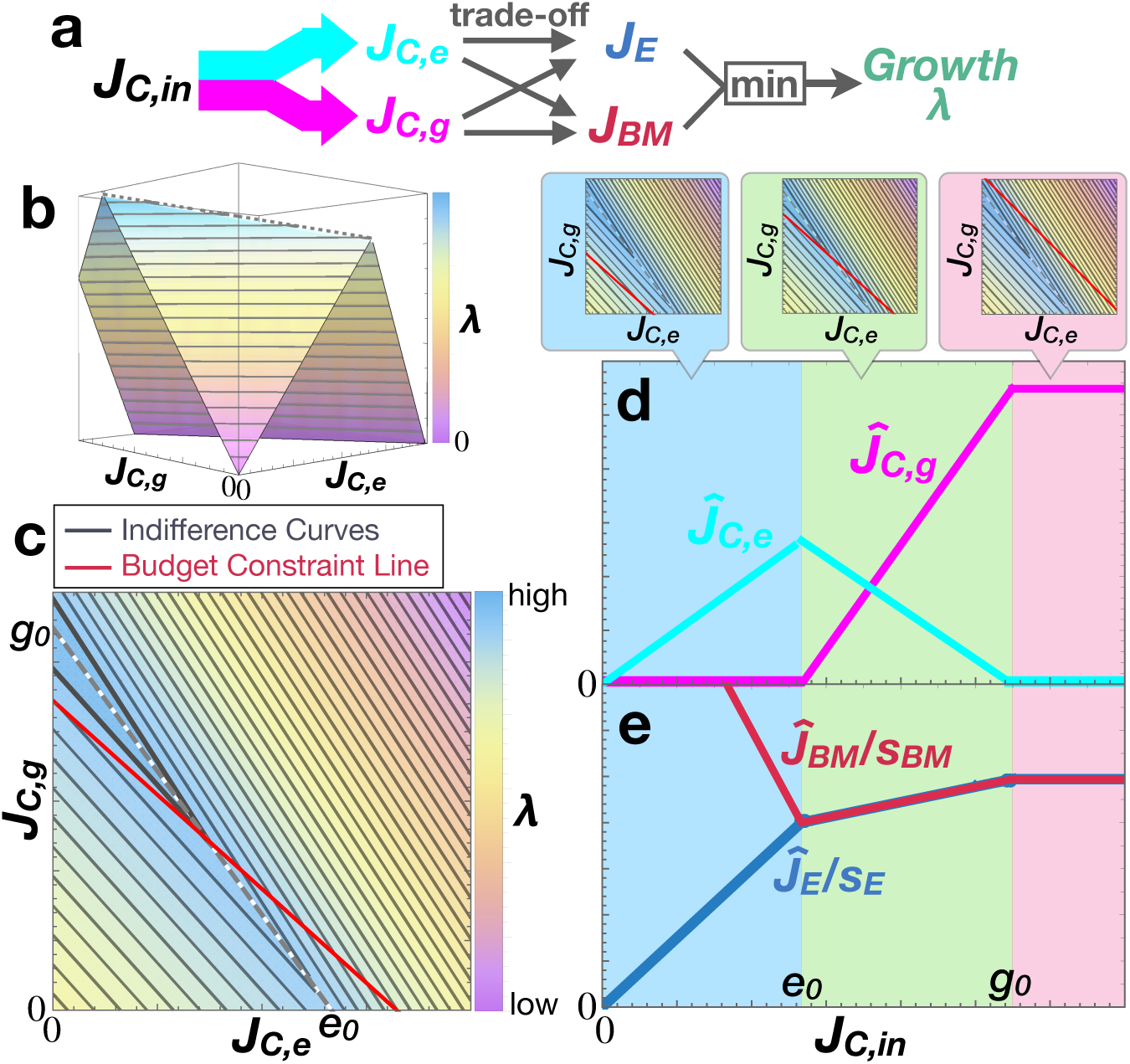
Overflow metabolism and Giffen behaviour as an optimization problem. **a**, Schematic illustration of the microeconomics model for overflow metabolism. **b**, Landscape of the growth rate λ(*J*_*C,e*_,*J*_*C,g*_). **c**, Contour map of the growth rate. Contours of the growth rate (i.e., indifference curves) and the budget constraint line (Eq. (1)) are represented by black and red solid lines, respectively. The grey dashed line is the ridge-line of the objective function (Eq. (2)). The background colour represents the growth rate λ(*J*_*C,e*_,*J*_*C,g*_). **d**, Dependence of the optimal strategy (*Ĵ*_*C,e*_, *Ĵ*_*C,g*_) on *J*_*C,in*_ (Engel curve; Eq. [S7] in Supplementary Note 4). *J*_*C,e*_ (cyan line) and *J*_*C,g*_ (magenta line) in the optimal solutions are plotted against *J*_*C,in*_. **e**, *Ĵ*_*E /*_ *s*_*E*_ ≡ *J*_*BM*_(*Ĵ*_*C,e*_, *Ĵ*_*C,g*_)/*s*_*E*_ (blue line) and *Ĵ*_*B,M /*_ *s*_*BM*_ ≡ *J*_*BM*_(*Ĵ*_*C,e*_, *Ĵ*_*C,g*_)/*s*_*BM*_ (dark-red line) in the optimal solutions are plotted against *J*_*C,in*_. The blue line also corresponds to the optimized growth rate λ(*Ĵ*_*C,e*_, *Ĵ*_*C,g*_). The top panels depict the contour maps and the budget constraint line for the cases *J*_*C,in*_ ≤ *e*_0_ (light-blue area), *g*_0_ ≥ *J*_*C,in*_ ≥ *e*_0_ (light-green area), and *J*_*C,in*_ ≥ *g*_0_ (pink area).

The cellular growth rate, λ(*J*_*C,e*_, *J*_*C,g*_), which is given as a function of the metabolic fluxes, is the utility (objective function) for a cell. Because cells have to build their components from precursors of biomass and energy molecules for successful division, λ(*J*_*C,e*_, *J*_*C,g*_) is determined from the production rates of energy molecules *J*_*E*_(*J*_*C,e*_, *J*_*C,g*_) and biomass precursors *J*_*BM*_ (*J*_*C,e*_, *J*_*C,g*_). The energy production flux, *J*_*E*_, is the sum of the fluxes of ATP synthesis via oxidative phosphorylation, *J*_*E,e*_, and aerobic glycolysis, *J*_*E,g*_, which are proportional to *J*_*C,e*_ and *J*_*C,g*_ with different coefficients, respectively:

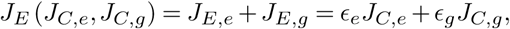

where *ϵ*_*e*_ and *ϵ*_*g*_ are the efficiency of metabolism determined from stoichiometry, and satisfy *ϵ*_*e*_ *> ϵ*_*g*_ *>* 0 because oxidative phosphorylation produces a greater amount of ATP than glycolysis from the same amount of the carbon source.

In contrast, by denoting the total amount of a given resource other than carbon (e.g., the intracellular space in the solvent capacity hypothesis (15, 16)) by *ρ*_tot_, the competition for the limited resource is described as *ρ*_*e*_ + *ρ*_*g*_ + *ρ*_*BM*_ = *ρ*_tot_, where *ρ*_*e*_, *ρ*_*g*_, and *ρ*_*BM*_ are the resources allocated to oxidative phosphorylation, aerobic glycolysis, and production of biomass precursors, respectively. Based on the law of mass action, each flux is proportional to the allocated resource in the steady state: 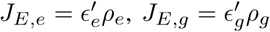, and 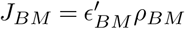. It follows that

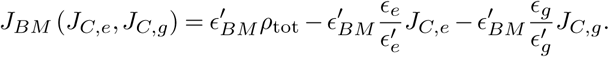

Empirical observations show that oxidative phosphorylation requires more resources than glycolysis with lactate secretion, and thus 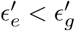 holds (see Supplementary Note 3 for estimation of the parameters). Consequently, there is a trade-off between *J*_*E*_(*J*_*C,e*_, *J*_*C,g*_) and *J*_*BM*_ (*J*_*C,e*_, *J*_*C,g*_).

Here, each cellular component including biomass, is built by metabolic reactions following the rules of stoichiometry. In general, due to the law of mass conservation, the compounds involved in biochemical reactions cannot be replaced by each other. Accordingly, the total amount of the product is determined by the least abundant component. This property of stoichiometry is identical to the concept of (perfect) complementarity in microeconomics, which is represented by a Leontief utility function, i.e., the least available element of all goods (17). Hence, the growth rate is represented by a Leontief utility function as

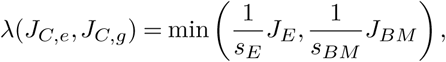

where *s*_*E*_ and *s*_*BM*_ are the stoichiometric coefficients for energy molecules and biomass precursors to synthesize biomass respectively (Fig. 1b). Then, the contour (i.e., indifference curve in microeconomics) is given as a two-valued function, as shown in Fig. 1c (Eq. [S7] in Supplementary Information). The growth rate is maximized at the tangent point of the budget constraint line (Eq. (1)) to the contour with the largest growth rate.

We first show that the response of overflow metabolism against the intake of carbon is explained as a result of optimization. As shown in Fig. 1d, the dependence of the optimal carbon allocation (*Ĵ*_*C,e*_, *Ĵ*_*C,g*_) on the income *J*_*C,in*_, called the Engel curve in microeconomics, is calculated in the case without inefficiency in metabolism, i.e., *p*_*e*_ = *p*_*g*_ =1 (Eq. [S8] in Supplementary Information). This Engel curve and the dependence of the optimal growth rate on *J*_*C,in*_ (the blue curve in Fig. 1e) are in good agreement with experimental observations (14–16, 25, 27).

If the influx of carbon sources *J*_*C,in*_ is lower than 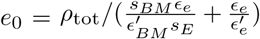, the budget constraint line has no tangent point to any contour, and the maximum growth rate is achieved at (*J*_*C,e*_, *J*_*C,g*_)= (*J*_*C,in*_, 0) (see the light-blue area in Figs. 1d and 1e). In this regime, the occupancy of limited resources *ρ*_tot_ does not limit cell growth, and all the carbon intake is used for oxidative phosphorylation to produce energy molecules more efficiently.

When the carbon intake *J*_*C,in*_ is sufficiently high, the budget constraint line has a tangent point to a contour. The set of such tangent points with various *J*_*C,in*_ is given as the line on which 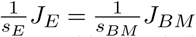 (see the ridgeline in Figs. 1b and 1c, represented by the dashed lines):

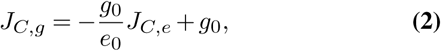

where 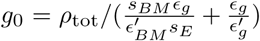. Accordingly, the growth rate is maximized at the intersection between the above ridge-line (2) and the budget constraint line (1). Because the slope of the ridgeline (2) is negative due to the trade-off between the efficiency of energy production and resource occupancy, when *J*_*C,in*_ increases, the growth rate λ(*J*_*C,e*_, *J*_*C,g*_) and *J*_*C,g*_ increases and *J*_*C,e*_ decreases along the ridgeline (see the light-green area in Figs. 1d and 1e); i.e., overflow metabolism occurs. From the viewpoint of microeconomics, this indicates a negative income effect. Because the Leontief utility function has a null substitution effect (28, 29), overflow metabolism is immediately identified as Giffen behaviour in which the electron transport chain or respiratory pathway is a Giffen good.

When *J*_*C,in*_ increases further and exceeds *g*_0_, the optimal solution always takes the value (*Ĵ*_*C,e*_, *Ĵ*_*C,g*_) = (0,*g*_0_) (see the pink area in Figs. 1d and 1e) because the global maximum of the growth rate is achieved at this point. In this situation, the efficiency of producing energy molecules is no longer a primary concern given the excess carbon sources available, and the occupancy of the limited resources is the only selection criterion for the two metabolic pathways.

Notably, if there is no trade-off, i.e., both *ϵ*_*e*_ *> ϵ*_*g*_ and 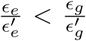 hold, the maximal point of the growth rate is not (*J*_*C,e*_, *J*_*C,g*_)= (0, *g*_0_) but rather (*J*_*C,e*_, *J*_*C,g*_)= (*e*_0_, 0). Then, cells allocate the carbon intake to only the electron transport chain until reaching the upper bound determined by the available resources at *J*_*C,e*_ = *e*_0_, whereas *J*_*C,g*_ remains at zero; that is, overflow metabolism cannot be observed. A trade-off is thus required for overflow metabolism and Giffen behaviour.

Based on the correspondence between overflow metabolism and Giffen behaviour, our theory can explain the mechanism of the inverse Warburg effect (19–22, 30). In Giffen behaviour, the demand for a Giffen good increases with its price. In metabolism, when uncouplers of oxidative phosphorylation are administered, it becomes less efficient due to dissipation of the proton gradient to produce ATP (31), i.e., the price of the electron transport chain *p*_*e*_ increases. Then, the carbon flux toward the electron transport chain will counterintuitively increase (Fig. 2). This seemingly wasteful behaviour is indeed observed ubiquitously, e.g., in yeasts (32, 33) and fungi (34). Moreover, addition of such drugs is known to induce hyperthermia (23, 31). Giffen behaviour also connects drug-associated overheating with overflow metabolism by the same mechanism: drug administration increases the flow in the more dissipative oxidative phosphorylation.

**Fig. 2.**
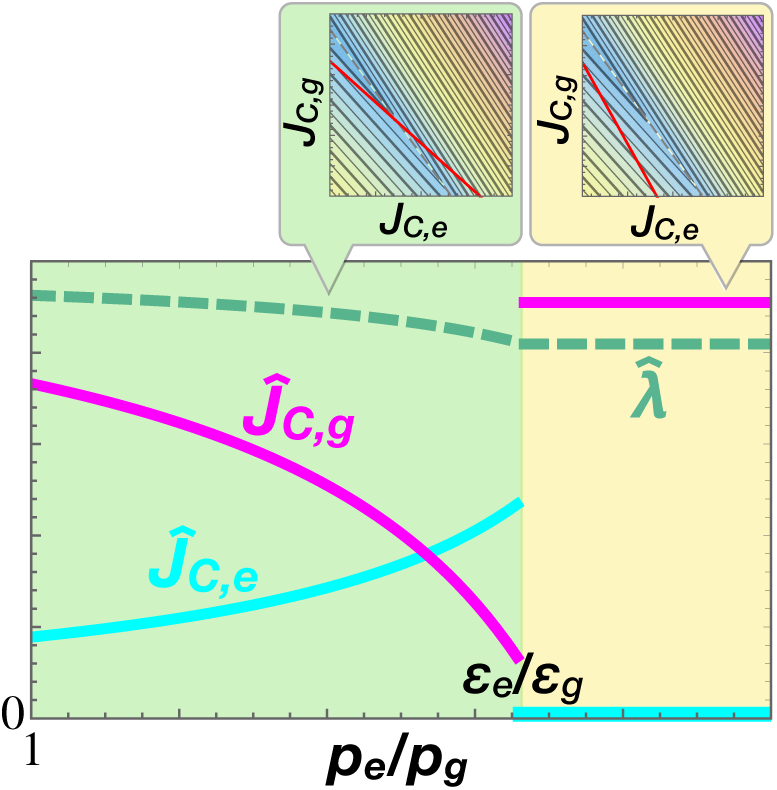
Dependence of the optimal strategy on the price of the electron transport chain 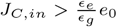 and *p*_*g*_ =1 are fixed here. The cyan, magenta, and green curves depict *Ĵ*_*C,e*_, *Ĵ*_*C,g*_, and 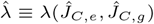, respectively (scaled with different units). The top panels depict the contour maps and the budget constraint lines for the regimes (I) *p*_*e*_ *< ϵ*_*e*_*/ϵ*_*g*_ (light-green area) and (II) *p*_*e*_ *> ϵ*_*e*_*/ϵ*_*g*_ (yellow area).

Of note, the property of optimal carbon allocation (*Ĵ*_*C,e*_, *Ĵ*_*C,g*_) qualitatively switches depending on whether the price ratio *p*_*e*_*/p*_*g*_ is higher or lower than *ϵ*_*e*_*/ ϵ*_*g*_. In regime (I) *p*_*e*_*/p*_*g*_ *< ϵ*_*e*_*/ ϵ*_*g*_ (the light-green area in Fig. 2), the demand *J*_*C,e*_ increases along with the price *p*_*e*_, i.e., Giffen behaviour is observed. In contrast, in regime (II) *p*_*e*_*/p*_*g*_ *> ϵ*_*e*_*/ ϵ*_*g*_ (the yellow area in Fig. 2), using only glycolysis as (*J*_*C,e*_, *J*_*C,g*_)= (0, *J*_*C,in*_*/p*_*g*_) is optimal in terms of growth rate. In this regime, considering the imbalance in price, the glycolytic pathway becomes more efficient for both *J*_*E*_ and *J*_*BM*_, and thus there is no longer a trade-off (Supplementary Note 2). As shown in Fig. 2, the optimal allocation of carbon fluxes discontinuously jumps from regime (I) to (II) when *p*_*e*_ increases, whereas the optimal growth rate changes continuously. In addition, in the case of *ϵ*_*e*_*e*_0_*/ ϵ*_*g*_ *> J*_*C,in*_ *> e*_0_, *Ĵ*_*C,g*_ reaches zero at *p*_*e*_*/p*_*g*_ = *J*_*C,in*_*/e*_0_ and then *Ĵ*_*C,e*_ continuously decreases before switching from regime (I) to (II) (Supplementary Figure 1). In fact, sudden stimulation of fermentation and reduction of respiration against the addition of uncouplers above a critical concentration has been observed experimentally (see Figure 2 in ref. (33)).

Our study was based on the spirit inherited from constraint-based modelling in that evolutionary processes force metabolic systems to become optimized (1–4), but we adopted a reductionist approach here. By constructing a simplified utility landscape comprising only a few variables with the aid of microeconomics, we uncovered the minimal, universal structure for overflow metabolism, which no longer depends on the specific assumption of limited resource allocation (Supplementary Note 2). Hence, besides overflow metabolism, a variety of metabolic systems are expected to show Giffen behaviour as long as they satisfy the two requirements of trade-off and complementarity. Here, perfect complementarity necessarily holds in metabolic systems owing to the law of mass conservation, i.e., complementarity represents the requirement of balance among metabolic fluxes for optimization. Hence, Giffen behaviour will be general for metabolic systems with a trade-off such as the Embden-Meyerhoff-Parnass and Entner-Doudoroff glycolysis pathways (35, 36) and the mixed-acid or lactic-acid fermentation pathways (37). Notably, same as overflow metabolism, if the flux of a metabolic pathway drops against the increased substrate intake, the flux is predicted to increase by making the metabolic pathway less efficient. The optimal solutions on the ridgeline in our theory can reproduce the phenomenological model in ref. (25), though it is not explicitly formulated as an optimization problem and thus, cannot systematically explain the metabolic strategies in response to drugs. Therefore, it could not address, for example, the link between the Warburg effect and the inverse Warburg effect.

From the perspective of microeconomics, we provide a concrete example of and a novel prediction for Giffen goods. The results are consistent with and contribute toward resolving the mechanism for an economic experiment (18) in which Giffen behaviour is observed with moderate income, but cannot be observed in the case of extreme poverty. In our model, Giffen behaviour occurs in the intermediate range of income at which the trade-off matters (Fig. 1), whereas a good behaves as the perfect substitute for the other one when the income is sufficiently low. Note that perfect complementarity does not necessarily exist for Giffen behaviour outside of metabolism (Supplementary Note 5).

We have paved the road for the field of microeconomics of metabolism, using overflow metabolism and Giffen behaviour as stepping stones. This will bring about further development in both biology and economics. For instance, since overflow metabolism is commonly characteristic to both cancer and stem cells, our theory will contribute to understanding the origin of cancer stem cells (38). We also expect that the metabolic behaviours characteristic to cancer such as serine and glycine metabolism (39, 40) and the metabolic responses to drugs (23) will be understood by the microeconomics of metabolism.

## Box 1 Theory of consumer choice in microeconomics

The theory of consumer choice explains how the price *p*_*i*_ of goods and income *I* determine consumption behaviour, i.e., the demand *x*_*i*_ for each good *i* under the given utility (i.e., objective function) *u*(*x*_1_, *x*_2_, …, *x*_*n*_). If the consumer is sufficiently rational in their decisions, the decision-making process is mapped onto an optimization problem of the utility under the budget constraint 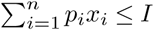. When the number of goods *n* is two, the utility landscape is represented as a curved surface, and the demand for each good is basically determined from a tangent point of the budget constraint line, *p*_1_*x*_1_ + *p*_2_*x*_2_ = *I*, to the indifference curve (i.e., contour of objective function) on which the utility takes the largest value. In the example of the utility function *u*(*x*_1_, *x*_2_) shown in the Box Figure, the demand for each good increases when its price decreases or income increases, and vice versa. Such goods are termed normal goods in microeconomics.

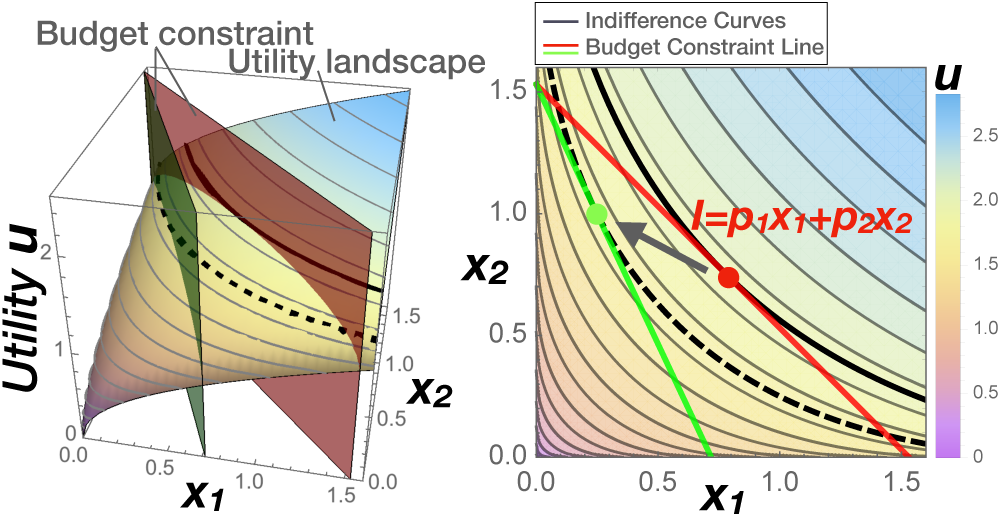

The influence of a change in price on the demand for goods can be decomposed into two distinct effects, known as the Slutsky equation (5, 6) (see Supplementary Note 1 for details):

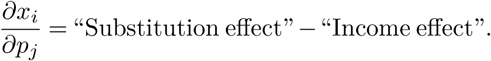

The substitution effect is caused by relative changes in the combination of the demand for goods. Since the self-substitution effect has to be non-negative (6), the demand for a good cannot be increased by the substitution effect when its price rises. In contrast, an increase in a good’s price effectively decreases the budget to spend freely. Such an effective change in income alters the demand for each good, which is called the income effect. The income effect of a good can be either positive or negative, by which the demand for the good increases or decreases with the income increasing, respectively (see Box Table). The goods with a negative income effect are termed inferior goods. A Giffen good, which is a particular type of inferior goods, shows a negative income effect that is larger than its self-substitution effect.

| Type of goods | Self-substitution effect | Income effect | Income ↑ | Price ↑ |
| --- | --- | --- | --- | --- |
| Normal good | non-positive | positive | Demand ↑ | Demand ↓ |
| Inferior good | non-positive | (slightly) negative | Demand ↓ | Demand ↓ |
| Giffen good | non-positive | negative | Demand ↓ | Demand ↑ |

## ACKNOWLEDGMENTS

We would like to acknowledge Chikara Furusawa for helpful discussions and careful reading of the manuscript, and Kunihiko Kaneko, Atsushi Kamimura, and Yasushi Okada for useful comments.

## AUTHOR CONTRIBUTIONS

J.F.Y. and T.S.H. designed and performed the research, and wrote the paper.

