## Supplementary Information for "Microeconomics of metabolism: Overflow metabolism as Giffen behaviour"

### Supplementary Note 1. Substitution Effect and Income Effect

The influence of a change in price on the demand for goods can be decomposed into an income effect and a substitution effect as mentioned in the main text. This is known as Hicksian decomposition or the Slutsky equation (1).

We here define $x_i(\mathbf{p}, I)$ as the (optimal) demand for good $i$ as a function of the price of goods $\mathbf{p}$ and income $I$. $E(\mathbf{p}, u)$ represents the minimum income required for a given utility value $u$ under price $\mathbf{p}$, and thus $h_i(\mathbf{p}, u) \equiv x_i(\mathbf{p}, E(\mathbf{p}, u))$ is the smallest demand for good $i$ necessary to achieve the given utility value $u$. We can differentiate this with respect to $p_j$ and obtain

$$\frac{\partial h_i(\mathbf{p}, \hat{u}(\mathbf{p}, I))}{\partial p_j} = \frac{\partial x_i(\mathbf{p}, I)}{\partial p_j} + \frac{\partial x_i(\mathbf{p}, I)}{\partial I} \frac{\partial E(\mathbf{p}, \hat{u}(\mathbf{p}, I))}{\partial p_j},$$

where $\hat{u}(\mathbf{p}, I)$ is the maximal utility with the given price $\mathbf{p}$ and income $I$. Note that the last term, $\partial E(\mathbf{p}, u(\mathbf{p}, I))/\partial p_j$, is simply equal to the demand $x_j$ under price $\mathbf{p}$ and income $I$.

Accordingly, the change in $x_i(\mathbf{p}, I)$ due to a change in $p_j$ is given by the Slutsky equation:

$$\frac{\partial x_i(\mathbf{p}, I)}{\partial p_j} = \frac{\partial h_i(\mathbf{p}, \hat{u}(\mathbf{p}, I))}{\partial p_j} - x_j(\mathbf{p}, I) \frac{\partial x_i(\mathbf{p}, I)}{\partial I}. \quad \text{[S1]}$$

The first term, $\partial h_i(\mathbf{p}, u)/\partial p_j$, is the substitution effect, which is caused by relative changes in the combination of price values (2). The case of $i = j$ represents the self-substitution effect, which is proven to be always non-positive (1), i.e., the substitution effect never increases the demand for a good when its own price increases. In contrast, the second term, $x_j(\mathbf{p}, I) \frac{\partial x_i(\mathbf{p}, I)}{\partial I}$, is the income effect, which can be either positive or negative. This effect reflects the change in the demand for goods due to the effective decrease of income that is caused by increasing the price of a good. According to Eq. [S1], these two effects determine the dependence of the demand for goods on the price (see also the Box Table in the main text).

The substitution effect is represented as movement of the combination of the demand for goods along an indifference curve to a point at which the tangent line has a slope that is equal to the ratio of the altered price of goods. Hence, if the utility is given as a Leontief utility function, the substitution effect is zero within a certain range of the price change due to the indifferentiability of each indifference curve at the kink. It follows that the substitution effect of the utility is zero in the case of metabolic systems. Consequently, whether or not a metabolic pathway is a Giffen good depends only on the sign of its income effect; namely, a metabolic pathway behaves as a Giffen good if its income effect is negative. In the case of overflow metabolism, when the budget constraint line intersects with the ridgeline (Eq. [2] in the main text), the income effect is negative for the utility $\lambda(J_{C,e}, J_{C,g})$, and thus Giffen behaviour is observed.

Note that although the demand for branded goods also increases with their price, they are not Giffen goods. This is because the demand for branded goods increases with the income, in other words, their income effect is positive. Such goods are called Veblen goods in microeconomics (3, 4).

### Supplementary Note 2. The theory of consumer choice for overflow metabolism

The general theory for two goods and two complementary "objectives" elucidates the minimal requirements for Giffen behaviour, which is applicable to a variety of phenomena in biology and economics.

Let us define a Leontief utility function

$$u(x_1, x_2) \equiv \min(A, B), \quad \text{[S2]}$$

where two complementary objectives, $A$ and $B$, are defined as

$$\begin{cases} A(x_1, x_2) = a_1 x_1 + a_2 x_2 + A_0, \\ B(x_1, x_2) = b_1 x_1 + b_2 x_2 + B_0. \end{cases}$$

The demand for and price of two goods are represented by $x_1, x_2$ and $p_1, p_2$, respectively. The budget constraint line is thus $I = p_1 x_1 + p_2 x_2$.

With respect to utility [S2], the model proposed for overflow metabolism in the main text corresponds to the case where the signs of the parameters are given as $\mathrm{sgn}\begin{pmatrix} a_1 & a_2 \\ b_1 & b_2 \end{pmatrix} = \begin{pmatrix} + & + \\ - & - \end{pmatrix}$, while other sets of parameters also evoke overflow metabolism and Giffen behaviour. For instance, a trade-off between the efficiencies of producing different molecules such as ATP and NADPH (5) can also cause overflow metabolism, where the parameters are given by $\mathrm{sgn}\begin{pmatrix} a_1 & a_2 \\ b_1 & b_2 \end{pmatrix} = \begin{pmatrix} + & + \\ + & + \end{pmatrix}$ (Supplementary Figure 2). Indeed, the Leontief utility function with $\mathrm{sgn}\begin{pmatrix} a_1 & a_2 \\ b_1 & b_2 \end{pmatrix} = \begin{pmatrix} + & + \\ + & 0 \end{pmatrix}$ was recently reported to demonstrate Giffen behaviour in the context of microeconomics (6, 7), which is a special case of the utility function [S2].

In this section, we demonstrate that the optimization problem of the above utility [S2] with every set of parameters will always be reduced to the optimization problem with an identical structure to that shown in the main text, as long as it shows Giffen behaviour. Hence, the two conditions, i.e., complementarity and trade-off, are required in all cases. Moreover, we expand the trade-off to include the effect of price.

First, we consider the simple case where $p_1 = p_2 = 1$. We here assume a trade-off between goods 1 and 2 as $a_1 > a_2$ and $b_1 < b_2$, whereas without the trade-off, the optimal strategy is to use either good 1 or 2 only. Owing to the trade-off, the utility is maximized on the ridgeline

$$x_2 = -\frac{a_1 - b_1}{a_2 - b_2}x_1 - \frac{A_0 - B_0}{a_2 - b_2} \qquad [S3]$$

where $A(x_1, x_2) = B(x_1, x_2)$ holds. Giffen behaviour can be observed when this ridgeline has a negative slope and exists on the first quadrant, i.e., when $-(a_1 - b_1)/(a_2 - b_2) < 0$ and $(B_0 - A_0)/(a_2 - b_2) > 0$ hold.

From symmetry between $A(x_1, x_2)$ and $B(x_1, x_2)$, it is sufficient to consider the situation where

$$a_1 > b_1, \quad a_2 > b_2, \quad B_0 > A_0 \qquad [S4]$$

are satisfied. Note that this is the condition for a comparative advantage (8).

If both $b_1$ and $b_2$ are non-positive and $a_1$ and $a_2$ are positive, the condition [S4] is autonomously satisfied. Then, the income effect is negative and Giffen behaviour is observed, as shown in the main text (see Fig. 1).

Even if $b_1$ or $b_2$ is positive, the condition [S4] can be satisfied as long as the condition $\mathrm{sgn}(a_1 - b_1) = \mathrm{sgn}(a_2 - b_2)$ is satisfied, as in Supplementary Figure 2. As discussed below, even such cases can be reduced to an optimization problem with the same universal structure demonstrated in the main text.

When Giffen behaviour can occur (i.e., the condition [S4] is satisfied), we can take a real number $c$ such that $a_2 > c > b_2$, and the utility [S2] can be represented as

$$\begin{aligned} u(x_1, x_2) = \min(A, B) &= c(x_1 + x_2) + A_0 + \min\left((a_1 - c)x_1 + (a_2 - c)x_2, (b_1 - c)x_1 + (b_2 - c)x_2 + B_0 - A_0\right) \\ &= cI + A_0 + \min(A', B'), \end{aligned} \qquad [S5]$$

where

$$\begin{cases} A'(x_1, x_2) = (a_1 - c)x_1 + (a_2 - c)x_2 = a_1'x_1 + a_2'x_2, \\ B'(x_1, x_2) = (b_1 - c)x_1 + (b_2 - c)x_2 + B_0 - A_0 = b_1'x_1 + b_2'x_2 + B_0'. \end{cases}$$

Here, $a_1' = a_1 - c > a_2 - c = a_2'$, $b_1' = b_1 - c < b_2 - c = b_2' < 0$, and $B_0' = B_0 - A_0 > 0$ are satisfied from the condition [S4]. Because only the last term, $\min(A', B')$, depends on allocation of the income to goods, the generalized model is reduced to the same optimization problem as proposed in the main text as long as it shows Giffen behaviour.

A similar argument remains valid even when considering changes in price, although in this case the definition of trade-offs needs to be expanded to include the effect of price. In this case, a real number $c$ can be taken such that $a_2/p_2 > c > b_2/p_2$. Then, the utility [S2] can be represented as

$$\begin{aligned} u(x_1, x_2) &= c(p_1x_1 + p_2x_2) + A_0 + \min\left((a_1 - cp_1)x_1 + (a_2 - cp_2)x_2, (b_1 - cp_1)x_1 + (b_2 - cp_2)x_2 + B_0 - A_0\right) \\ &= cI + A_0 + \min(A', B'), \end{aligned} \qquad [S6]$$

where

$$\begin{cases} A'(x_1, x_2) = (a_1 - cp_1)x_1 + (a_2 - cp_2)x_2 = a_1'x_1 + a_2'x_2, \\ B'(x_1, x_2) = (b_1 - cp_1)x_1 + (b_2 - cp_2)x_2 + B_0 - A_0 = b_1'x_1 + b_2'x_2 + B_0'. \end{cases}$$

If $a_1/a_2 > p_1/p_2 > b_1/b_2$ holds, $a_1' = a_1 - cp_1 > a_2 - cp_2 = a_2'$, $b_1' = b_1 - cp_1 < b_2 - cp_2 = b_2' < 0$, and $B_0' = B_0 - A_0 > 0$ are satisfied. Again, the generalized models showing Giffen behaviour are reduced to the optimization problem with the same universal structure as demonstrated in the main text.

Of note, if the difference of the price between goods 1 and 2 is so large that $p_1/p_2$ is larger than $a_1/a_2$ or smaller than $b_1/b_2$, Giffen behaviour disappears even when the condition [S4] and the trade-off of efficiencies to produce $A$ and $B$ (i.e., $a_1 > a_2$ and $b_1 < b_2$) hold. In this case, we need to consider a trade-off including the price. If $p_1/p_2$ is larger than $a_1/a_2$, a unit of demand for good 2 produces more $A$ than that for good 1 even when $a_1$ is higher than $a_2$. That is, there is no longer a trade-off between the production of $A$ and $B$. Hence, the condition for a trade-off is rewritten as $a_1/p_1 > a_2/p_2$ and $b_1/p_1 < b_2/p_2$ (or $a_1/p_1 < a_2/p_2$ and $b_1/p_1 > b_2/p_2$).

Intuitively, the trade-off including the price corresponds to rescaling of the utility landscape so that the slope of the budget constraint line, $-\frac{p_1}{p_2}$, becomes equal to $-1$. If there is a trade-off between $A$ and $B$ in the rescaled landscape (i.e., $a_1/p_1 > a_2/p_2$ and $b_1/p_1 < b_2/p_2$, or $a_1/p_1 < a_2/p_2$ and $b_1/p_1 > b_2/p_2$), Giffen behaviour can be observed.

### Supplementary Note 3. Estimation of parameters

In the main text, we used an arbitrarily chosen set of parameters ($\epsilon_e = 0.75, \epsilon_g = 0.45, \epsilon_e' = 0.5, \epsilon_g' = 0.7, \rho_{tot} = 0.6, \epsilon_{BM}' = 1, s_E = s_{BM} = 0.5$) in order to demonstrate a mathematical structure of the landscape clearly (see also Supplementary Table 1 for meaning of the symbols). However, our results do not depend on the precise values of the parameters as long as the relative values of parameters satisfy the conditions described in the main text: $\epsilon_e > \epsilon_g$ and $\epsilon_e' < \epsilon_g'$.

One can estimate the parameter set in actual cells and show that the estimated parameters indeed satisfy the conditions for overflow metabolism. The details are given below (see also Supplementary Table 2).

First, the intake rate of the carbon source (e.g., glucose) is the order of 1 [mMol glucose/gDW] (9).

According to stoichiometry, mitochondrial respiration can generate 36 ATP molecules per 1 molecule of glucose at most, while in reality it generates about $30-32$ ATP molecules per 1 molecule of glucose (10, 11). In contrast, aerobic glycolysis generates 2 ATP and 4 NADH, in total, from 1 glucose, and fermentation generates 2 ATP more. Besides, in the aerobic environment, 1 NADH molecule can be converted into, at most, 2.5 ATP molecules; thus, at most, 12 or 14 ATP molecules can be generated from 1 glucose by aerobic glycolysis or fermentation, respectively (10, 12). Then, the values of $\epsilon_e$ and $\epsilon_g$ are dependent on the organism in question, but a requirement for overflow metabolism, $\epsilon_e > \epsilon_g$, is always satisfied: $\epsilon_e \simeq 32$ [mMol ATP/ mMol glucose] and $\epsilon_g = 2 \sim 12$ [mMol ATP/ mMol glucose]. Notably, the decrease in the optimal flux $\hat{J}_{C,e}$ against nutrient supply $J_{C,in}$ can be very gentle (whereas glycolysis $\hat{J}_{C,g}$ increases steeply), e.g., in the situation where $\epsilon_e/\epsilon_g$ is relatively large (i.e., NADH produced in glycolysis is not converted to ATP consuming oxygen), as experimentally observed (11, 13).

$\rho_{tot}$, $\epsilon_e'$, and $\epsilon_g'$ depend on the nature of the limited resource in question. When the limited resource $\rho_{tot}$ is the total volume of mitochondria or solvent capacity of mitochondria in cancer cells, muscle cells, yeasts, et al.:the intracellular volume for proteins per cellular dry weight is $\rho_{tot} \simeq 3 \times 10^{-3}$ [L/gDW] and the approximate values of the other parameters are $\epsilon_e' \simeq 3 \times 10^3$ [mMol ATP/L/hour], $\epsilon_g' \simeq 9 \times 10^4$ [mMol ATP/L/hour], and $s_{BM}/\epsilon_{BM}' \simeq 10^{-3}$ [hour × L/gDW] (11, 14). Alternatively, when $\rho_{tot}$ is the fraction of enzymes for growth in E. coli: the approximate values of the parameters are estimated as $\rho_{tot} \simeq 2 \times 10^{-1}$, $\epsilon_e' \simeq 4 \times 10^2$ [mMol ATP/gDW/hour] and $\epsilon_g' \simeq 8 \times 10^2$ [mMol ATP/gDW/hour], and $s_{BM}/\epsilon_{BM}' \simeq 10^{-1}$ [hour] (12).

When the balance between cellular redox state and energy demand is considered (15), $\rho_{tot}$ is the maximal flux of $NAD^+$ produced by the other cellular processes and is estimated as $\rho_{tot} \simeq 3 \times 10^2$ [mMol $NAD^+$/gWW/hour]. Then, $\rho_g = J_{E,g}/\epsilon_g'$ is 0 (i.e., $\epsilon_g'$ and $\epsilon_e'/\epsilon_g'$ can be regarded as infinity and zero, respectively) because glycolysis does not consume $NAD^+$. In contrast, respiration consumes 1 $NAD^+$ per 1 glucose, or per $\epsilon_e$ ATP molecules; thus, $\epsilon_e'$ equals $\epsilon_e$[mMol ATP/mMol $NAD^+$]. In addition, the amount of required $NAD^+$ per cellular biomass is estimated as $s_{BM}/\epsilon_{BM}' \simeq 2 \times 10^2$ [mMol $NAD^+$/gWW].

Again, the other requirement for overflow metabolism, $\epsilon_e' < \epsilon_g'$, is satisfied in all the above cases.

Finally, $p_e$ and $p_g$ approximately equal 1 without the administration of drugs because, at most, only $\sim 10\%$ carbon is usually leaked as intermediates of central metabolism (16). In contrast, under the existence of drug such as uncouplers of respiration, $p_e$ is substantially larger than 1, quantified as the inverse of the ATP yield, i.e., the fraction of the non-dissipated proton.

### Supplementary Note 4. Dependence of the optimal strategy on the income and price

The growth rate $\lambda(J_{C,e}, J_{C,g})$ takes the value $\tilde{\lambda}$, the contour (i.e., indifference curve in microeconomics) is given as a two-valued function, as shown in Fig. 1c in the main text:

$$J_{C,g} = \begin{cases} -\frac{\epsilon_e}{\epsilon_g} J_{C,e} + \frac{s_E}{\epsilon_g}\tilde{\lambda}, & \text{if } \frac{1}{s_E} J_E \le \frac{1}{s_{BM}} J_{BM} \\ -\frac{\epsilon_e}{\epsilon_g}\frac{\epsilon_g'}{\epsilon_e'} J_{C,e} + \frac{\epsilon_g'}{\epsilon_g}\left(\rho_{\text{tot}} - \frac{s_{BM}}{\epsilon_{BM}'}\tilde{\lambda}\right). & \text{if } \frac{1}{s_E} J_E \ge \frac{1}{s_{BM}} J_{BM} \end{cases} \quad \text{[S7]}$$

The growth rate is maximized at the tangent point of the budget constraint line (Eq. [1]) to the contour with the largest growth rate.

In the case of $p_e = p_g = 1$, the dependence of the optimal strategy $(\hat{J}_{C,e}, \hat{J}_{C,g})$ on $J_{C,in}$, called the Engel curve in microeconomics, is calculated (see also Fig. 1c in the main text):

$$\left(\hat{J}_{C,e}, \hat{J}_{C,g}\right) = \begin{cases} (J_{C,in}, 0)\,, & \text{if } J_{C,in} \le e_0 \\ \left(\frac{e_0}{g_0 - e_0}(g_0 - J_{C,in}), \frac{g_0}{g_0 - e_0}(J_{C,in} - e_0)\right), & \text{if } g_0 \ge J_{C,in} \ge e_0 \\ (0, g_0)\,. & \text{if } J_{C,in} \ge g_0 \end{cases} \quad \text{[S8]}$$

Changes in price alter the demand for goods, and thus the optimal strategy depends on the price $p_e$ and $p_g$ as well as on the income $J_{C,in}$. Accordingly, if the price $p_e$ or $p_g$ takes a value larger than 1, the Engel curve is generalized as follows.

$$\left(\hat{J}_{C,e}, \hat{J}_{C,g}\right) = \begin{cases} (J_{C,in}/p_e, 0)\,, & \text{if } J_{C,in} \le p_e e_0 \\ \left(\left(g_0 - \frac{J_{C,in}}{p_g}\right) \Big/ \left(\frac{g_0}{e_0} - \frac{p_e}{p_g}\right), \left(\frac{J_{C,in}}{p_e} - e_0\right) \Big/ \left(\frac{p_g}{p_e} - \frac{e_0}{g_0}\right)\right), & \text{if } p_g g_0 \ge J_{C,in} \ge p_e e_0 \\ (0, g_0)\,, & \text{if } J_{C,in} \ge p_g g_0 \end{cases} \quad \text{[S9]}$$

with $e_0 = \rho_{\text{tot}}/(\frac{s_{BM}\epsilon_e}{\epsilon_{BM}' s_E} + \frac{\epsilon_e}{\epsilon_e'})$ and $g_0 = \rho_{\text{tot}}/(\frac{s_{BM}\epsilon_g}{\epsilon_{BM}' s_E} + \frac{\epsilon_g}{\epsilon_g'})$. Here, the optimal $(J_{C,e}, J_{C,g})$ for the case $p_g g_0 \ge J_{C,in} \ge p_e e_0$ is obtained from the intersecting point of the budget constraint line with the ridgeline (Eq. [2] in the main text). Note that Eq. [S8] is identical to Eq. [S7] if $p_e = p_g = 1$ is satisfied.

### Supplementary Note 5. Giffen behaviour requires complementarity but not perfect complementarity

Based on biological considerations, we introduced the Leontief utility function $\lambda(J_{C,e}, J_{C,g})$ that has perfect complementarity in the main text. However, Giffen behaviour can be observed for utility functions with only partial complementarity, as long as the substitution effect is sufficiently small.

An example of such utility functions is

$$u(x_1, x_2) \equiv \left[ s_E / \left( \epsilon_e x_1 + \epsilon_g x_2 \right) + s_{BM} / \left( \rho_{\mathrm{tot}} - \frac{\epsilon_e}{\epsilon_e'} x_1 - \frac{\epsilon_g}{\epsilon_g'} x_2 \right) \epsilon_{BM}' \right]^{-1}. \quad \text{[S10]}$$

The landscape of this utility function (Supplementary Figure 3A) is similar to that of $\lambda(J_{C,e}, J_{C,g})$ (Fig. 1a in the main text). As shown in Supplementary Figure 3B, $x_1$ with the utility [S9] showing Giffen behaviour within a range of the price of the good 1, $p_1$.

Another example of utility functions showing Giffen behaviour is $u(J_{C,e}, J_{C,g}) \equiv J_E H(J_{BM} - J_{BM,min})$, where $H(\cdot)$ is the Heaviside step function.

**Supplementary Table 1. Biological and economic meanings of symbols.**

| Symbol | Biological meaning | Economic meaning |
|---|---|---|
| $J_{C,in}$ | Intake flux of carbon source | Income |
| $J_{C,e}, J_{C,g}$ | Fluxes of carbon in oxidative phosphorylation and glycolysis | Demand for goods |
| $p_e, p_g$ | Inefficiency of metabolism in oxidative phosphorylation and glycolysis | Price of goods |
| $\lambda$ | Growth rate | Utility |
| $J_E$ | Total flux of ATP production | An objective |
| $J_{E,e}, J_{E,g}$ | Flux of ATP production by oxidative phosphorylation and glycolysis | |
| $J_{BM}$ | Total flux of biomass precursors production | Another objective |
| $\rho_{\text{tot}}$ | Total amount of the limited resource | |
| $\rho_e, \rho_g, \rho_{BM}$ | Fraction of the limited resource used for oxidative phosphorylation, glycolysis, and biomass synthesis | |
| $\epsilon_e, \epsilon_g$ | Stoichiometric efficiency of ATP production in oxidative phosphorylation and glycolysis | |
| $\epsilon'_e, \epsilon'_g, \epsilon'_{BM}$ | Occupancy rate of the limited resource for oxidative phosphorylation, glycolysis, and biomass synthesis | |
| $s_E, s_{BM}$ | Stoichiometric constants | |

**Supplementary Table 2. Estimated parameters**

| Parameter | Value | Reference |
|---|---|---|
| $\epsilon_e$ | $\sim 30$ [mMol ATP/mMol glucose] | Refs. (10) |
| $\epsilon_g$ | $2 \sim 14$ [mMol ATP/mMol glucose] | Refs. (10) |
| $p_e$ | $\gtrsim 1$ | Ref. (16) |
| $p_g$ | $\sim 1$ | Ref. (16) |
| $s_E$ | $6 \times 10^1$ [mMol ATP/gDW] | Ref. (17) |
| Solvent capacity hypothesis | | Ref. (11, 14) |
| $\rho_{\text{tot}}$ | $3 \times 10^{-3}$ [L/gDW] | |
| $\epsilon'_e$ | $2 \times 10^4$ [mMol ATP/L/hour] | |
| $\epsilon'_g$ | $9 \times 10^4$ [mMol ATP/L/hour] | |
| $s_{BM}/\epsilon'_{BM}$ | $10^{-3}$ [hour $\times$ L/gDW] | |
| Proteome allocation hypothesis | | Ref. (12) |
| $\rho_{\text{tot}}$ | $2 \times 10^{-1}$ | |
| $\epsilon'_e$ | $4 \times 10^2$ [mMol ATP/gDW/hour] | |
| $\epsilon'_g$ | $8 \times 10^2$ [mMol ATP/gDW/hour] | |
| $s_{BM}/\epsilon'_{BM}$ | $10^{-1}$ [hour] | |
| Redox balance hypothesis | | Ref. (15) |
| $\rho_{\text{tot}}$ | $3 \times 10^2$ [mMol $NAD^+$/gWW/hour] | |
| $\epsilon'_e$ | $\epsilon_e$ [mMol ATP/mMol $NAD^+$] | |
| $\rho_g(\propto 1/\epsilon'_g)$ | 0 | |
| $s_{BM}/\epsilon'_{BM}$ | $2 \times 10^2$ [mMol $NAD^+$/gWW] | |

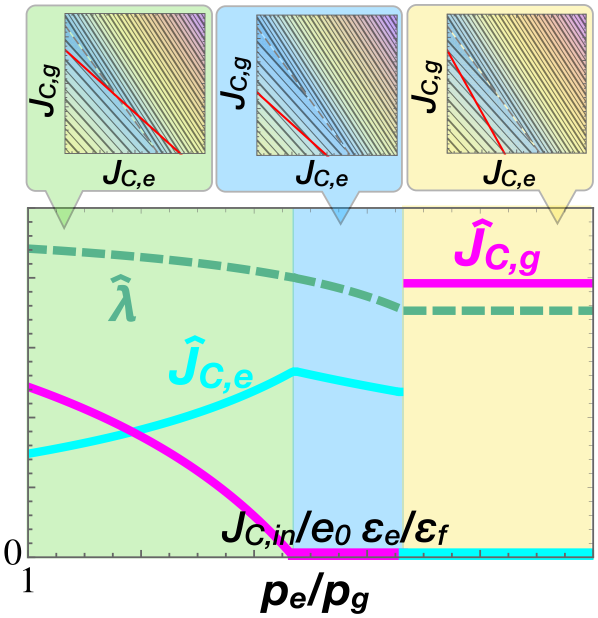


**Supplementary Figure 1.** Dependence of the optimal strategy $(\hat{J}_{C,e}, \hat{J}_{C,g})$ on the price of electron transport chain $p_e$ with $J_{C,in} < \frac{\epsilon_e}{\epsilon_g} e_0$. $J_{C,in} > e_0$ and $p_g = 1$ are fixed here. The cyan, magenta, and green curves depict $\hat{J}_{C,e}$, $\hat{J}_{C,g}$, and $\hat{\lambda} \equiv \lambda(\hat{J}_{C,e}, \hat{J}_{C,g})$, respectively (scaled with different units). Top panels depict the contour maps and the budget constraint lines for regime (I) $p_e < \epsilon_e/\epsilon_g$ (light-green area) and $J_{C,in} \leq p_e e_0$ and $p_e < \epsilon_e/\epsilon_g$ (light-blue area) and (II) $p_e > \epsilon_e/\epsilon_g$ (yellow area).

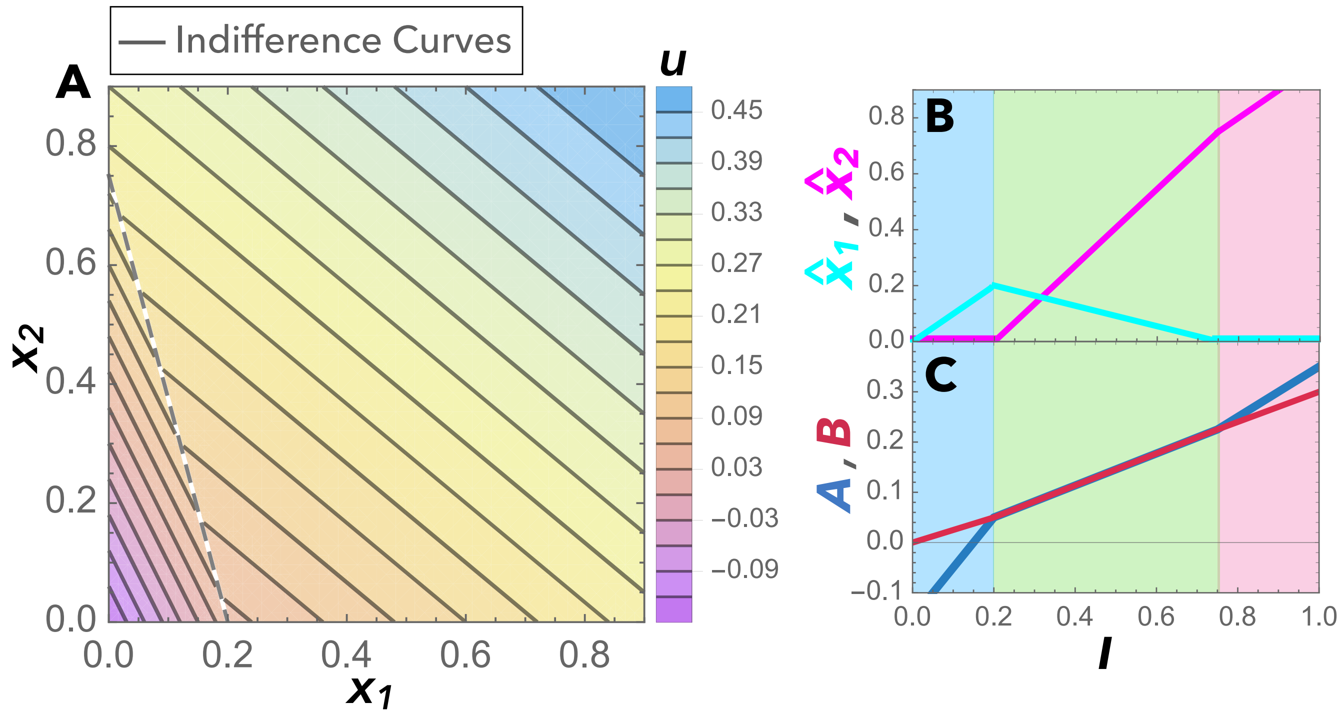


**Supplementary Figure 2.** Example of a generalized model with the coefficients $a_1, a_2, b_1, b_2 > 0$. (A) Contour map. Indifference curves are shown as grey lines, and the dashed line depicts the ridgeline [S3] on which $A(x_1, x_2) = B(x_1, x_2)$ holds. The background colour exhibits the value of the Leontief-type utility $u(x_1, x_2) = \min(A, B) = \min(a_1 x_1 + a_2 x_2 + A_0, b_1 x_1 + b_2 x_2 + B_0)$ (Eq. [S2]). (B) The Engel curve: $x_1$ (cyan line) and $x_2$ (magenta line) in the optimal solutions are plotted against the income $I$. (C) $A$ (blue line) and $B$ (dark-red line) in the optimal solutions are plotted against the income $I$. In (B-C), the price $p_1$ and $p_2$ are set at unity. In this example, the coefficients are set such that $a_1 > a_2 > b_2 > b_1$ and $B_0 > A_0$: $a_1 = 1, a_2 = 0.5, b_1 = 0.25, b_2 = 0.3, A_0 = -0.15, B_0 = 0$. Since there is a tradeoff between the production efficiencies of $A$ and $B$ and the condition [S4] is satisfied, Giffen behaviour is observed.

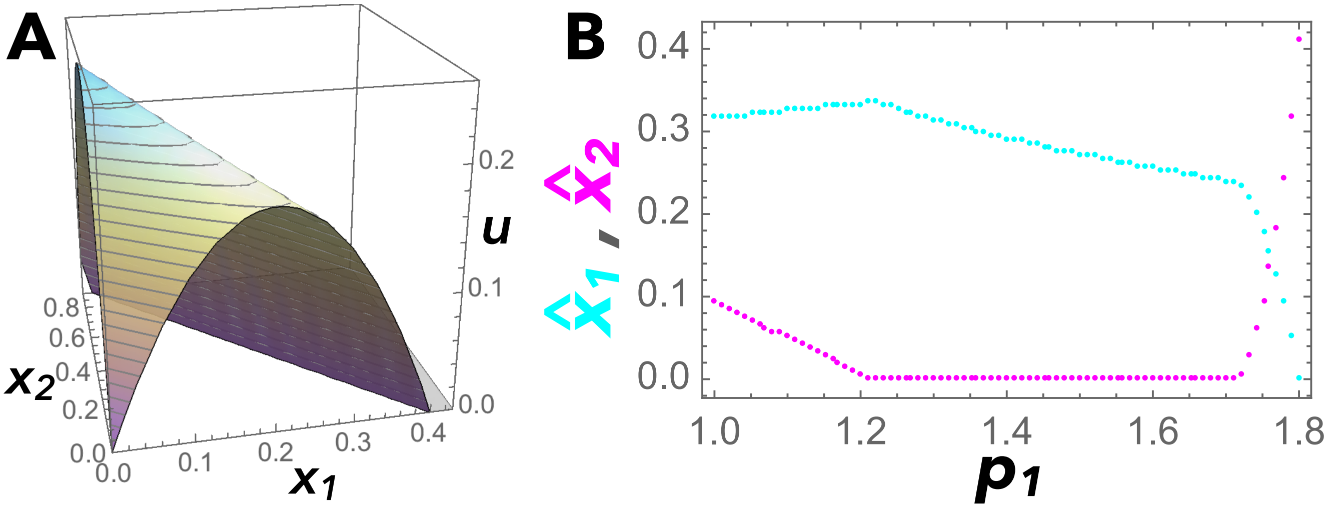


**Supplementary Figure 3.** Example of utility functions showing Giffen behaviour without perfect complementarity. The utility function $u(x_1, x_2)$ is given by Eq. [S9]. (A) Utility landscape of $u(x_1, x_2)$ and (B) demand curves for goods $1$ (light-blue dots) and $2$ (pink dots). The optimal strategies $(\hat{x}_1, \hat{x}_2)$ were numerically calculated with the parameters $\epsilon_e = 0.75, \epsilon_g = 0.4, \epsilon'_e = 0.5, \epsilon'_g = 1, J_{C,in} = 0.4$; all other parameters (i.e., $\rho_{\text{tot}}$, $\epsilon'_{BM}$, $s_E$, $s_{BM}$, and $p_2$) were set at unity.
